# DIRECTION-AWARE INTEGRATED BIOINFORMATICS ANALYSIS REVEALS CONCORDANT AND DISCORDANT MOLECULAR SIGNATURES LINKING TYPE 2 DIABETES MELLITUS AND POLYCYSTIC OVARY SYNDROME

**DOI:** 10.64898/2026.09.20.753005

**Authors:** Sodiq Okikiola Abdulsalam, Damilare Alabi Akanbi, Temiloluwa Esther Aloba, Merylin Wuraola Ogunlola, David Ifeoluwa Owopetu

**Affiliations:** Department of Biochemistry, University of Ibadan, Ibadan, Oyo State, Nigeria; Department of Physiology, University of Ibadan, Ibadan, Oyo State, Nigeria; Department of Zoology, University of Ibadan, Ibadan, Oyo State, Nigeria; Department of Microbiology, University of Ibadan, Ibadan, Oyo State, Nigeria

**Author notes:** Corresponding Author Name: Sodiq Okikiola Abdulsalam, Affiliation: Department of Biochemistry, University of Ibadan, Ibadan, Oyo State, Nigeria.

## Abstract

Polycystic ovarian syndrome and Type 2 diabetes mellitus are complex multivariate diseases that share well-established clinical pathophysiology, yet the molecular basis of their overlap is unknown. Prior bioinformatics studies have identified common differentially expressed genes between these two conditions. However, they have not accounted for the directionality of the gene expression changes observed, potentially obscuring any biological distinctions they may have. This study applied a direction-aware approach to classify molecular signatures between PCOS and T2DM as concordant or discordant. Transcriptomic analysis of GEO datasets GSE138518 (ovarian granulosa tissue, PCOS) and GSE25724 (pancreatic islet tissue, T2DM) identified 225 and 1,302 DEGs, respectively. Venn diagram analysis showed that only three genes (SLC6A8, RGS4, and SORL1) were shared between PCOS and T2DM from the total of 1,527 genes, and all 3 genes were regulated in opposing directions. Disease gene retrieval from the Comparative Toxicogenomics Database, Online Mendelian Inheritance in Man database, and GeneCards showed 214 shared disease-associated genes, with 311 genes unique to PCOS and 496 genes unique to T2DM. Protein-protein interaction construction using STRING (version 12.0) identified 70 interacting nodes. CytoHubba analysis across 6 scoring methods identified 18 high-confidence hub genes, including INS, BCL2, MTOR, LEP, MFN2, and PIK3CD. Functional enrichment analysis identified biological processes including immune activation, apoptosis, and insulin signaling, which were confirmed as the main pathways in KEGG pathway analysis. These findings show that while PCOS and T2DM share limited DEGs, they share common pathogenic pathways.

## INTRODUCTION

Polycystic ovarian syndrome (PCOS) and Type 2 Diabetes Mellitus (T2DM) are complex, multivariate metabolic diseases that have significant effects on world health. While PCOS is a complex endocrine condition characterized by hyperandrogenism, ovulatory dysfunction, and polycystic ovarian morphology, T2DM is characterized by persistent hyperglycemia due to insulin resistance and increasing pancreatic β-cell dysfunction [1, 2]. Growing clinical and molecular data suggests a substantial pathophysiology overlap between the two illnesses, despite the fact that they have historically been investigated as separate diseases [3].

Insulin resistance is a key factor in the development and course of PCOS and T2DM [4]. Insulin resistance in PCOS increases ovarian androgen production, which leads to hyperandrogenism in addition to metabolic dysfunction [5]. As a result, women with PCOS have a significantly higher chance of acquiring T2DM and decreased glucose tolerance in later life [6]. In addition to insulin resistance, these disorders are linked to oxidative stress, mitochondrial dysfunction, chronic low-grade inflammation, and dysregulation of glucose and lipid metabolism [1]. These similarities clearly imply that both diseases are caused by similar molecular pathways.

Despite this connection, little is known about the specific molecular interactions and regulatory networks that connect PCOS and T2DM. Though they frequently fall short of capturing the intricate, systems-level interactions that characterize multifactorial disorders, traditional experimental research has yielded important insights into specific pathways [7]. Recent developments in high-throughput technologies, including RNA sequencing and microarrays, have made it possible to thoroughly profile gene expression patterns across many disease states [8]. These datasets provide a systems biology view of disease caused by identifying differentially expressed genes (DEGs), functional enrichment pathways, and protein–protein interaction (PPI) networks using bioinformatics analysis [9].

One effective method for identifying common molecular markers between linked disorders is integrated bioinformatics analysis. This method improves statistical robustness and makes it easier to identify important genes and pathways that might not be visible in individual analyses by merging datasets from many research situations [10]. Numerous studies have looked for similar DEGs and signaling pathways in T2DM and PCOS, offering preliminary insights into their molecular interaction [11].

In a recent study, Zhang *et al*. [12] used bioinformatics methods to find important genes and pathways linked to PCOS and T2DM. The biological interpretability of their findings may be limited by methodological restrictions, despite the fact that their study is a significant step toward comprehending the underlying molecular foundation of these disorders. In particular, their approach used statistical thresholds like fold change and significance values to identify DEGs within each illness. An overlap analysis was then performed to identify common genes. Nevertheless, the directionality of gene expression changes, that is, whether genes were upregulated or downregulated in each condition, was not taken into consideration by this method.

Previous research may oversimplify the molecular link between T2DM and PCOS and perhaps mask important biological differences by treating all overlapping DEGs similarly, regardless of their direction of change. The necessity for a more sophisticated analytical approach that takes gene expression directionality into account when identifying common molecular signatures is highlighted by this constraint [12].

In order to close this gap, the current work uses an improved integrative bioinformatics technique that systematically classifies DEGs according to their expression patterns in addition to identifying common DEGs between T2DM and PCOS. In particular, concordant upregulated, concordant downregulated, and discordant expression groups will be used to categorize overlapping genes. This stratification makes it easier to identify pathways that are either simultaneously activated, jointly inhibited, or differently regulated across the two circumstances, allowing for a more comprehensive understanding of common and unique molecular processes.

Additionally, this study attempts to offer a more thorough and biologically significant interpretation of the molecular interactions between T2DM and PCOS by combining functional enrichment analyses, protein–protein interaction network construction, and hub gene identification in each expression category. In addition to helping to better understand the systems-level connections between metabolic and endocrine problems, such an approach is anticipated to enhance the discovery of relevant biomarkers and treatment targets.

While previous bioinformatics studies have laid the groundwork for exploring the molecular connections between T2DM and PCOS, there remains a critical need for approaches that incorporate expression directionality and network-level insights. The present study seeks to fill this gap by applying an integrative and direction-aware analytical framework to uncover robust and biologically relevant shared molecular signatures linking these two disorders [13].

## METHODOLOGY

### OBTAINING GENE DATA

Two gene expression datasets were retrieved from the National Center for Biotechnology Information (NCBI) Gene Expression Omnibus (GEO) database, namely GSE25724, which comprises gene expression data from human pancreatic islets, including 6 samples from type 2 diabetes patients and 7 non-diabetic controls, and GSE138518, which consists of RNA-sequencing data from 6 human ovarian granulosa tissue samples, including 3 PCOS-free donor controls and 3 patients diagnosed with polycystic ovarian syndrome. The GEO database was searched separately for the phrases “type 2 diabetes” and “polycystic ovarian syndrome.”

Differential gene expression analysis was performed using Gene Expression Omnibus 2 R (GEO2R), an online gene expression analysis tool integrated within the GEO platform for comparing gene expression profiles between different sample groups. GEO2R is based on the GEOquery and limma packages in the R programming environment. The GEOquery package enables access to GEO datasets, while the limma (Linear Models for Microarray Data) package was used to calculate log fold change (logFC) values and determine the statistical significance of differential gene expression between disease and control samples.

Heat maps were generated using GEO2R to visualize the expression patterns of differentially expressed genes across the selected samples. The resulting differential expression data were then exported and processed using Python. The NumPy library was used to filter genes using predefined thresholds, an adjusted p-value < 0.05, and |log2FC| > 1 to obtain the upregulated and downregulated genes.

#### Disease-Associated Gene Retrieval and Identification of Candidate Gene

To provide a comprehensive overview of genes associated with each disease, known disease-associated genes for PCOS and T2DM were obtained from three well-established databases: the Comparative Toxicogenomics Database (CTD), the Online Mendelian Inheritance in Man (OMIM) database, and GeneCards. Using three databases rather than one helps ensure that important genes are not missed.

Venn diagrams were generated using Venny 2.0 to determine shared and unique gene sets between T2DM and PCOS. For the DEGs, concordant and discordant gene sets were identified in a direction-specific manner by comparing upregulated and downregulated DEGs across the two diseases. In addition, Venn diagrams were used to determine overlapping disease-associated genes between T2DM and PCOS independently of expression direction or statistical filtering.

The intersecting gene sets obtained included 3 discordantly expressed DEGs and 214 shared disease-associated genes. These candidate gene sets were subsequently used for downstream functional enrichment and network-based analyses.

### IDENTIFICATION OF PROTEIN-PROTEIN INTERACTION

Protein–protein interactions were analyzed using the STRING database (Search Tool for the Retrieval of Interacting Genes/Proteins) as a comprehensive bioinformatics resource. STRING is a meta-database that integrates and scores both known and predicted interactions derived from multiple sources, including direct (physical) associations as well as indirect (functional) relationships, thereby providing a holistic view of protein connectivity. The database gathers information from computational prediction methods, experimental data, and publicly available repositories, allowing the systematic investigation of molecular interaction networks and a better understanding of cellular processes.

### TOPOLOGICAL METRICES

Topological sorting analysis was performed using Cytoscape for directed acyclic graphs (DAGs) to establish the hierarchical organization of network components. In this approach, a linear ordering of nodes was generated such that for every directed edge (u, v), node *u* precedes node *v*. This ordering is essential for modeling precedence relationships, including biological signaling pathways and dependency structures within complex networks.

### CONSTRUCTION OF TARGET NETWORK AND IDENTIFICATION OF HUB NODES

A protein–protein interaction (PPI) network was constructed using interaction data retrieved from the STRING database. The resulting network was imported into Cytoscape for visualization and further analysis. To ensure reliability, interactions with a confidence score of ≥ 0.4 were retained for downstream analysis.

Key hub genes within the network were identified using the CytoHubba plugin in Cytoscape. Six topological metrics were applied, including Degree, Maximal Clique Centrality (MCC), Maximum Neighborhood Component (MNC), Betweenness, Closeness, and Bottleneck. Degree centrality was used to identify highly connected nodes, with genes having ≥ 10 interactions considered as significant hubs.

MCC was used to detect nodes within highly interconnected subnetworks, while the remaining metrics provided complementary ranking based on different centrality measures. Genes consistently ranked within the top 10% across the selected algorithms were considered potential hub genes. Default parameters of CytoHubba were applied unless otherwise specified. A ranked list of important genes for additional validation and interpretation was produced by these results.

### FUNCTIONAL ENRICHMENT ANALYSIS OF DIFFERENTIALLY EXPRESSED GENES UTILIZING GENE ONTOLOGY (GO) AND ASSOCIATED PATHWAYS

The biological significance of the identified candidate genes was investigated using functional enrichment analysis in ShinyGO (version 0.85.1). The final candidate gene set, comprising the 214 shared disease-associated genes and the 3 overlapping differentially expressed genes (DEGs), was submitted for Gene Ontology (GO) enrichment analysis. GO annotation was performed across the three categories of Biological Process (BP), Cellular Component (CC), and Molecular Function (MF) to identify significantly enriched functional terms associated with the shared molecular signatures of Type 2 Diabetes Mellitus (T2DM) and Polycystic Ovary Syndrome (PCOS).

In addition, Kyoto Encyclopedia of Genes and Genomes (KEGG) pathway enrichment analysis was conducted to identify significantly enriched signaling and metabolic pathways associated with the candidate genes. Enriched GO terms and pathways with a false discovery rate (FDR) < 0.05 were considered statistically significant and were used for downstream biological interpretation.

## RESULT

### GEO Dataset Retrieval and Differential Gene Expression Analysis

Two gene expression datasets were downloaded from the National Centre for Biotechnology Information (NCBI) GEO database. The T2DM dataset (GSE25724) contained 13 samples from human pancreatic islets, comprising 7 non-diabetic controls and 6 T2DM patients. The PCOS dataset (GSE138518) contained 6 RNA-sequencing samples from human ovarian granulosa tissue, comprising 3 healthy controls and 3 PCOS patients. Differential expression analysis was carried out on both datasets using an adjusted p-value < 0.05 and |log2FC| > 1 as cutoff criteria, giving 1,302 differentially expressed genes (DEGs) for T2DM and 225 DEGs for PCOS.

A volcano plot visualization of GSE138518 (Fig. 1A) showed that the PCOS DEGs were spread across both upregulated and downregulated groups. Some strongly downregulated genes had −log10(Padj) values above 6, while several upregulated genes showed large positive fold changes beyond a log2FC of 6, indicating strong directional changes in gene expression in ovarian granulosa tissue in PCOS. For GSE25724 (Fig. 1B), the T2DM volcano plot showed a broader and more evenly spread pattern, with significant genes in both directions reaching −log10(P-value) scores above 5. The wider spread along the fold-change axis compared to the PCOS dataset reflects the greater magnitude of gene expression changes observed in pancreatic islet tissue under diabetic conditions. Together, the two datasets produced 1,527 DEGs, which were taken forward for further analysis.

**Fig. 1A and 1B:**
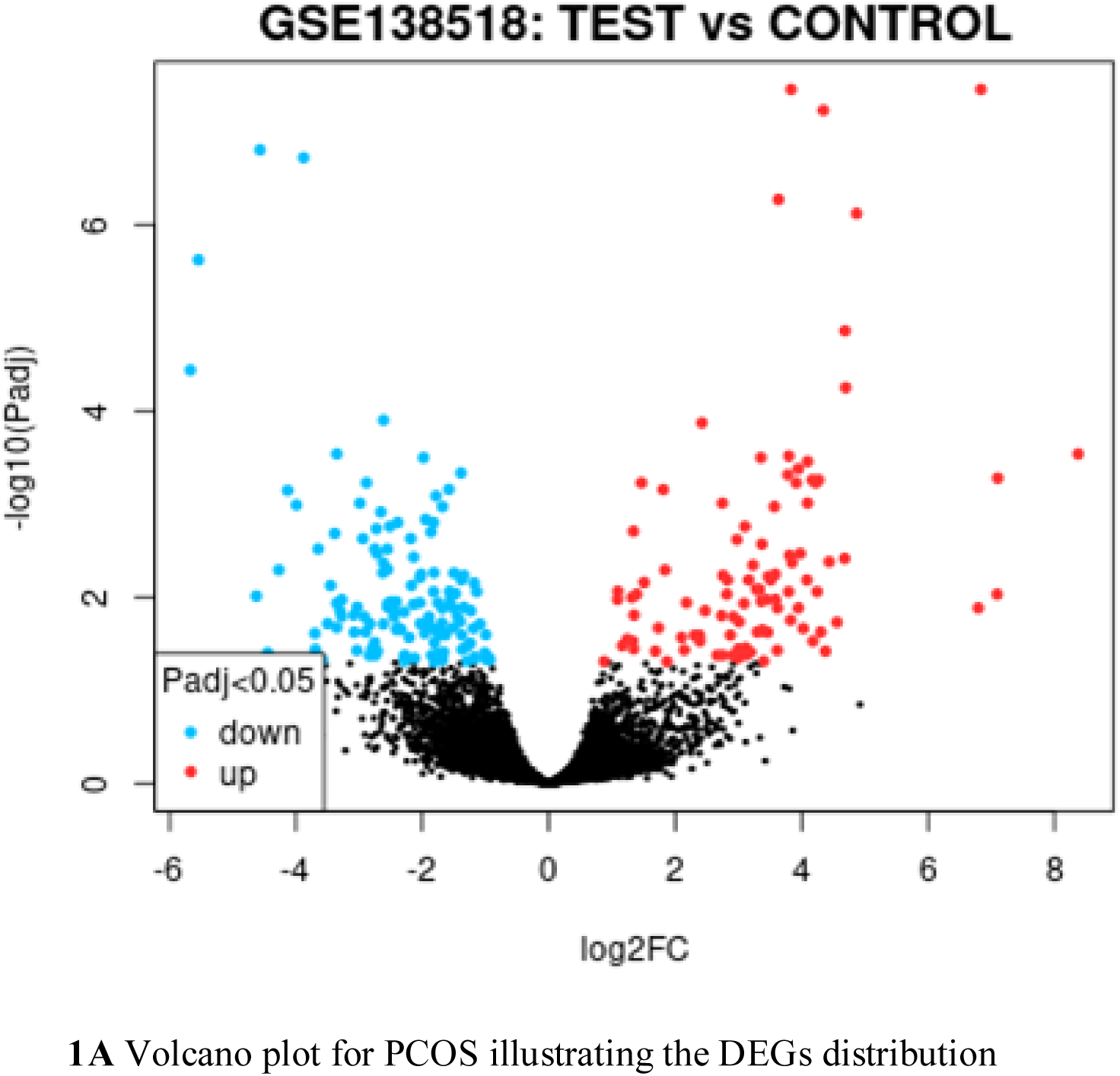
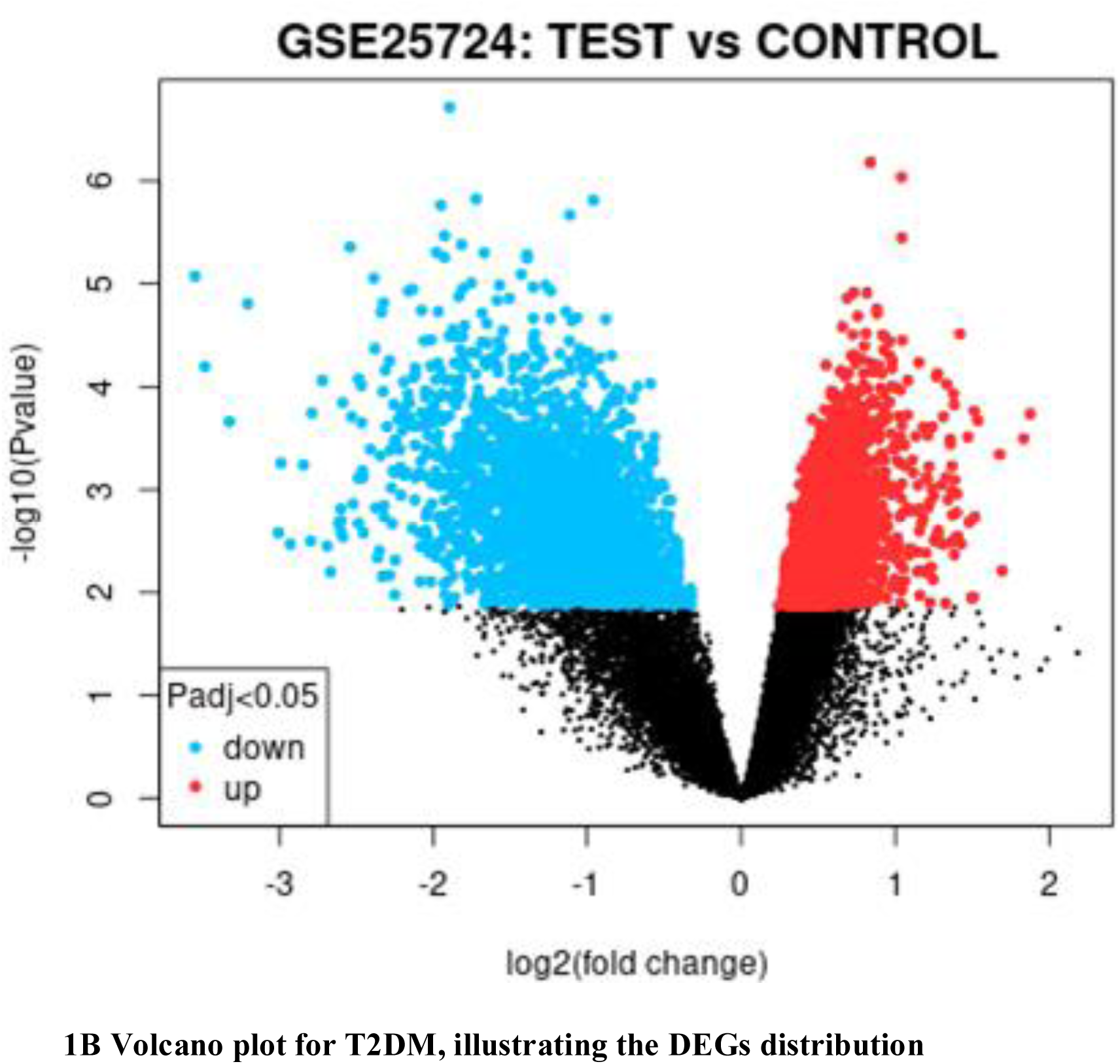
Volcano plots for PCOS and T2DM, respectively, were downloaded from GEO2R. Upregulated genes are represented in red, downregulated in blue, and non-significant genes in black.

**Table 1:** Summary Table of DEGs in T2DM and PCOS.

| **GEO Assession** | **Description** | **Total genes** | **Upregulated genes** | **Downregulated genes** |
| --- | --- | --- | --- | --- |
| **GSE25724** | Gene expression data from human pancreatic islets, consisting of 6 samples from type 2 diabetes patients and 7 non-diabetic controls | **1302** | **65** | **1237** |
| **GSE138518** | Comprises data from 6 human ovarian granulosa tissue samples, including 3 PCOS-FREE donor controls and 3 polycystic ovary syndrome disease patients. | **225** | **99** | **126** |

### Identification of Overlapping Differentially Expressed Genes

The DEG lists from both datasets were compared using a Venn diagram (Venny 2.1.0) to find genes that were differentially expressed in both conditions (Fig. 2A). Out of the 1,527 total DEGs, only 3 genes (0.2%) were shared between T2DM and PCOS, while 1,299 genes (85.2%) were unique to T2DM and 222 genes (14.6%) were unique to PCOS. This small overlap suggests that although both diseases show significant changes in gene expression, the specific genes affected are mostly different between the two conditions.

**Fig. 2A.**
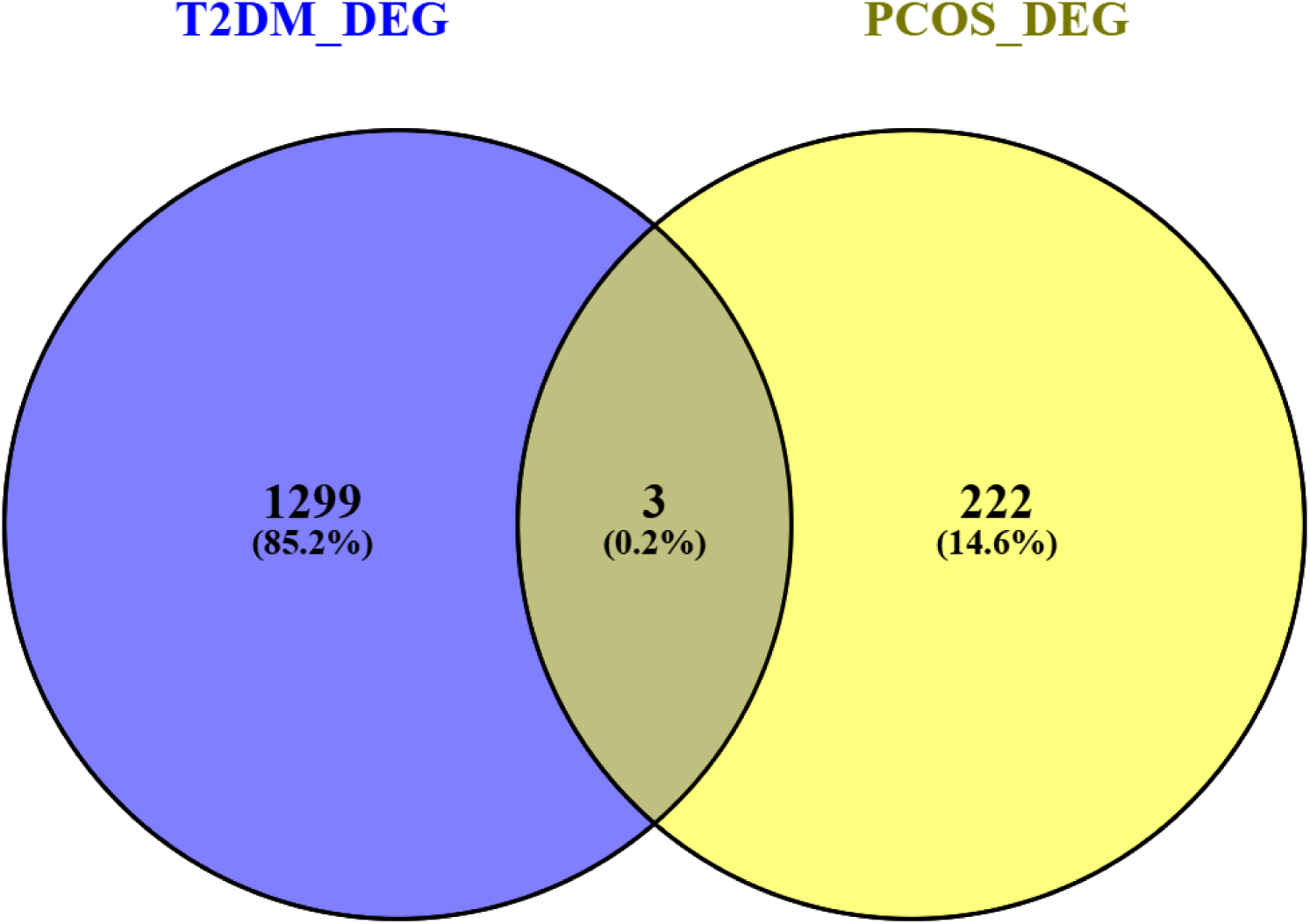
Overlapping genes between DEGs in T2DM and PCOS

A four-set Venn diagram was then used to compare the upregulated and downregulated gene lists from each disease separately (Fig. 2B). This showed that 1 gene (SLC6A8) was upregulated in T2DM but downregulated in PCOS, while 2 genes (RGS4 and SORL1) were upregulated in PCOS but downregulated in T2DM. All other intersections were empty, and no genes were concordantly upregulated or downregulated in both diseases.

**Fig. 2B.**
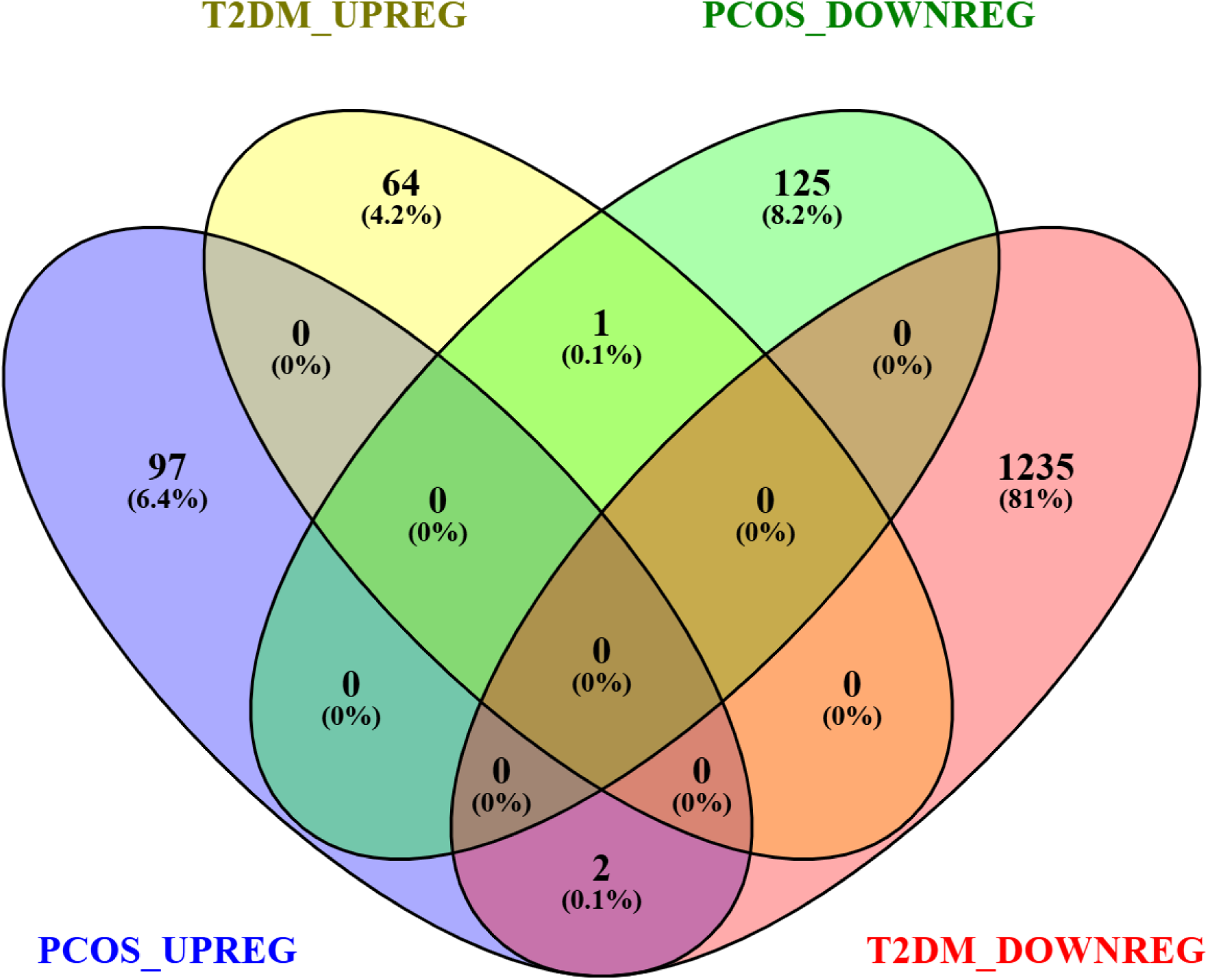
Four-set diagram to show the discordant relationship between the DEGs in T2DM and PCOS

The discordant regulation of all three shared genes is particularly notable. RGS4 and SORL1 are upregulated in PCOS but downregulated in T2DM, and SLC6A8 shows the opposite pattern, suggesting that even genes expressed in both conditions are regulated in opposing directions. This points to fundamentally different molecular responses in the two diseases rather than a shared activation or suppression of the same pathways, something that would be completely missed in a simple pooled gene overlap comparison.

### Disease Gene Retrieval and Candidate Gene List

Using three databases rather than one helps ensure that important genes are not missed. Using the three databases, the Comparative Toxicogenomics Database (CTD), the Online Mendelian Inheritance in Man (OMIM) database, and GeneCards, we got 525 disease genes for PCOS and 710 disease genes for T2DM. When the two lists were compared using a Venn diagram (Fig. 2C, 214 genes (21%) were shared between the two diseases, 311 genes (30.5%) were unique to PCOS, and 496 genes (48.6%) were unique to T2DM.

**Fig. 2C:**
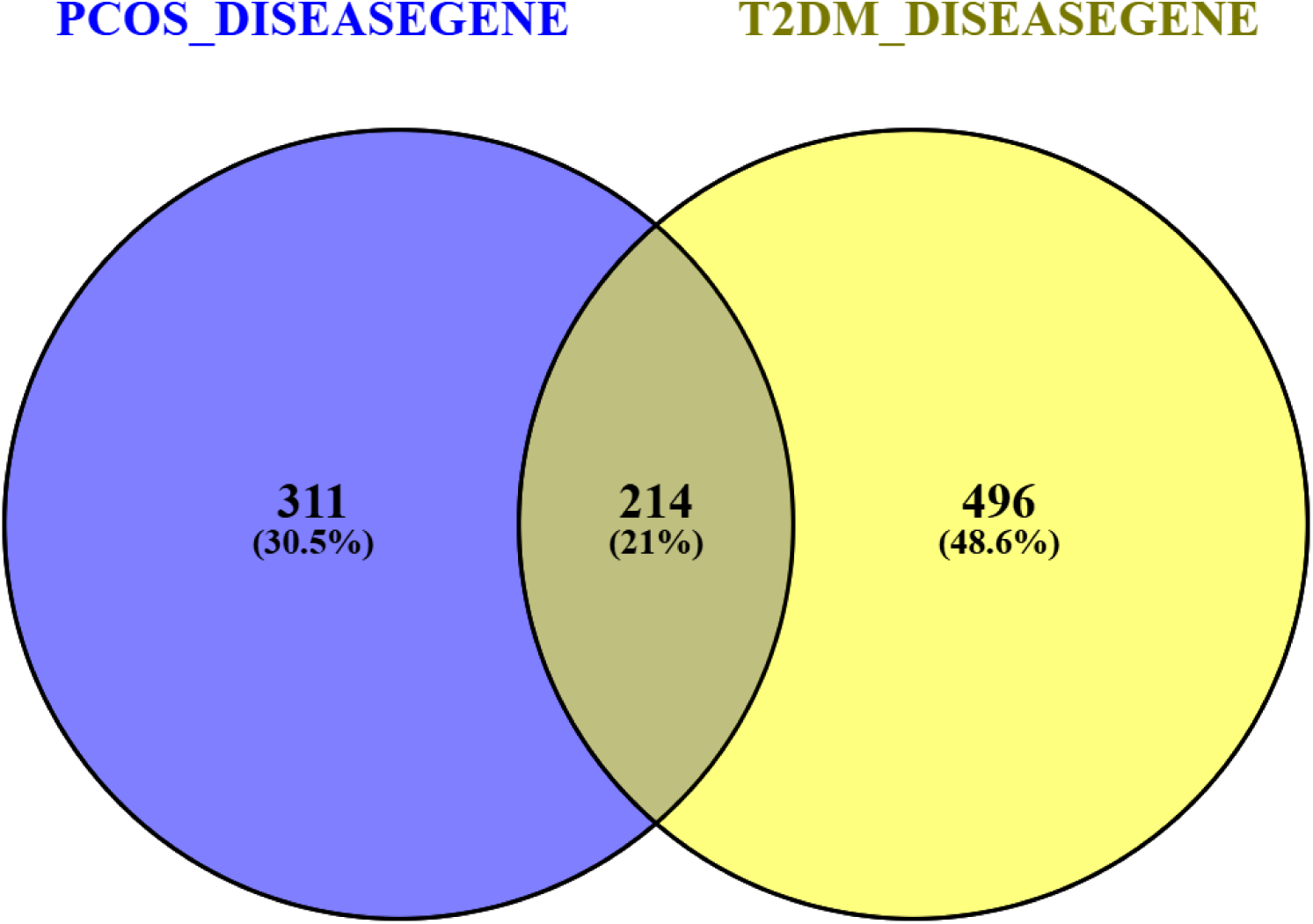
Overlapping genes between the disease genes gotten from CTD, OMIM, and GeneCards.

The fact that 214 genes are shared at the disease gene level, despite only 3 genes overlapping at the DEG level, suggests that while the two diseases do not show much similarity in their immediate gene expression responses, they do share a meaningful portion of their underlying genetic basis. This distinction is important and motivated the use of disease gene databases alongside expression data.

The final candidate gene list was curated by combining the 214 shared diseased genes with the 3 overlapping DEGs, giving a total of 217 candidate genes for downstream analysis.

### Protein–Protein Interaction Network Construction and Topological Analysis

The 217 candidate genes were entered into the STRING database (version 12.0) to build a protein-protein interaction (PPI) network, using a confidence score cutoff of 0.4. The network that was produced contained 70 interacting nodes and 64 edges (Fig. 3), meaning 70 of the 217 candidate proteins had at least one confirmed interaction with another protein in the network at the applied threshold.

**Fig. 3.**
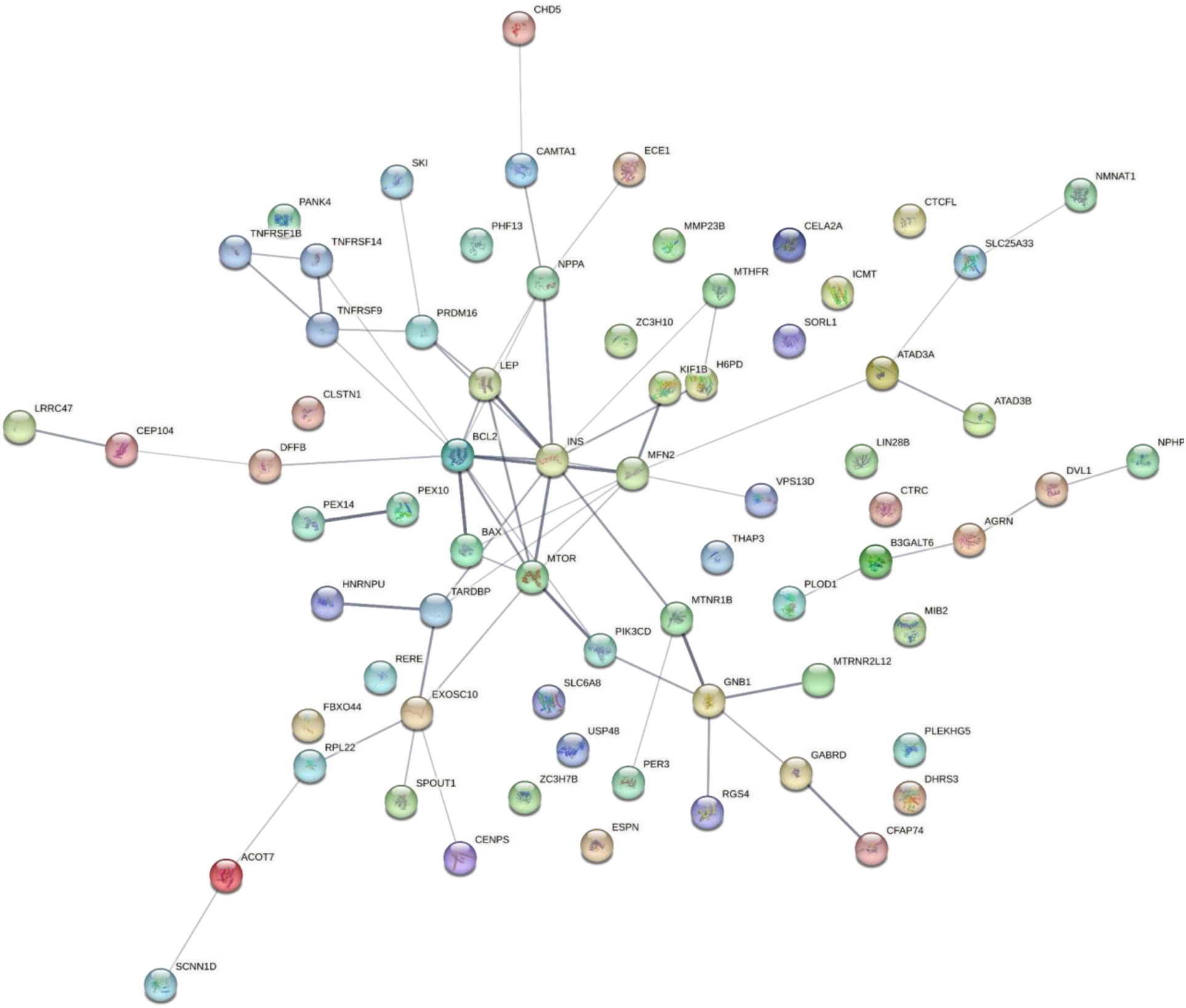
Protein–Protein Interaction Network Construction

The network showed significantly more interactions than would be expected by chance, with a PPI enrichment p-value of 0.0002. Since only 39 edges would be expected in a random set of proteins of the same size, the 64 observed edges show that the candidate proteins interact with each other far more than chance would predict, confirming that they form a biologically meaningful and connected network rather than a random collection of proteins.

The average number of interactions per protein (mean node degree) was 1.83, and the average local clustering coefficient was 0.395, meaning that proteins in the network tend to interact not just with each other but also with each other’s neighbors, forming tightly connected local groups.

Looking at the network visually, proteins such as INS, BCL2, MTOR, MFN2, and LEP stood out as having the highest number of connections, making them strong candidates for central regulatory roles in the shared biology of PCOS and T2DM.

### Hub Gene Identification by CytoHubba Analysis

The PPI network was imported into Cytoscape (Cytoscape_v3.10.4) and analyzed using CytoHubba to identify the most important hub genes. Six scoring methods were used: Degree, which counts how many direct connections each protein has; MCC (Maximal Clique Centrality), which identifies proteins sitting within the most densely connected parts of the network; EPC (Edge Percolated Component), which measures how much each protein contributes to the overall connectivity of the network; MNC (Maximum Neighborhood Component), which measures how densely connected a protein’s immediate neighbors are; Betweenness, which identifies proteins that act as key bridges controlling communication between different parts of the network; Bottleneck, which highlights critical proteins whose removal could disrupt network flow; and Closeness, which measures how efficiently a protein can interact with all other proteins in the network through the shortest paths. The top 20 genes were ranked by each method separately.

Genes that appeared consistently in the top rankings across all six methods were considered the most reliable hub genes (Table 2). A total of 18 genes were found to overlap across all six methods: INS, BCL2, MTOR, MFN2, LEP, BAX, PIK3CD, NPPA, GNB1, TARDBP, PRDM16, TNFRSF9, TNFRSF14, MTNR1B, MTHFR, H6PD, EXOSC10, and ATAD3A.

**Table 2:** Genes identified in the top rankings across all 6 methods, and identified as the most reliable Hub genes.

| RANK | DEGREE | MCC | MNC | CLOSENES<br>S | BETWEENN<br>ESS | BOTTLENE<br>CK |
| --- | --- | --- | --- | --- | --- | --- |
| 1 | BCL2 | LEP | LEP | INS | BCL2 | BCL2 |
| 2 | INS | BAX | PRMD16 | BCL2 | INS | INS |
| 3 | MFN2 | NPPA | BAX | MTOR | MFN2 | MFN2 |
| 4 | MTOR | EXOSC10 | NPPA | MFN2 | EXOSC10 | MTOR |
| 5 | LEP | MTOR | TNFRS14 | LEP | MTOR | EXOSC10 |
| 6 | NPPA | MFN2 | MFN2 | TARDBP | GNB1 | PIK3CD |
| 7 | EXCOSC10 | TNFRSF9 | MTOR | NPPA | NPPA | GNB1 |
| 8 | GNB1 | BCL2 | INS | PIK3CD | ATAD3A | AGRN |
| 9 | PRDM16 | GNB1 | TNFRSF9 | EXOSC10 | PIK3CD | NPPA |
| 10 | TNFRSF9 | INS | BCL2 | BAX | MTNR1B | ATAD3A |

The consistent identification of all 18 genes across six methodologically different algorithms strongly supports their reliability as true hub genes rather than method-specific results. These genes are therefore considered the high-confidence hub genes of the shared PCOS and T2DM network.

**Fig. 4.**
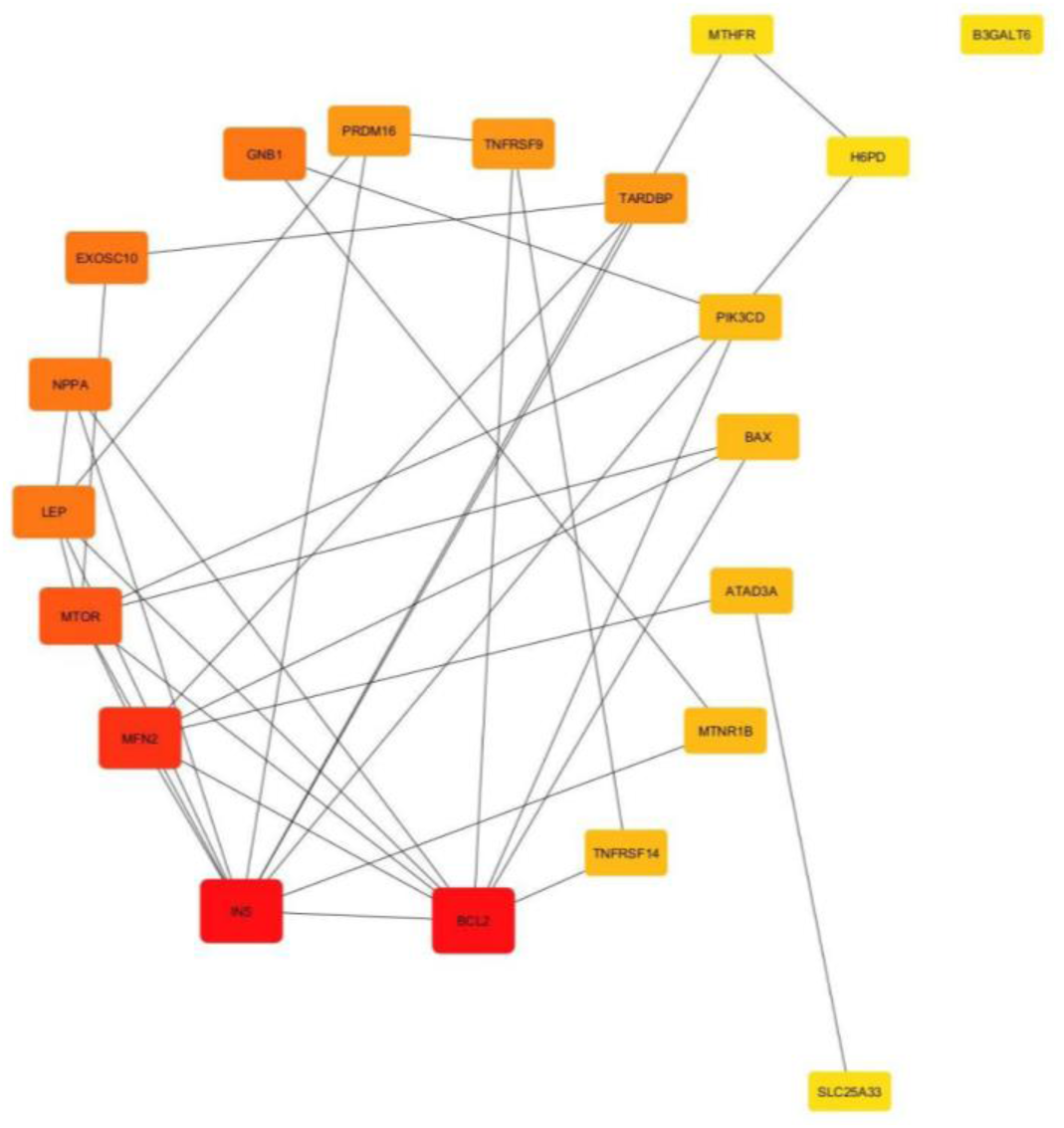
shows the PPI-network of the identified shared elements and the top 20 ranking of the nodes indicating the HUB genes using degree as ranking.

### Functional Enrichment Analysis

The 217 candidate genes were entered into ShinyGO (version 0.85.1) to run gene-set enrichment analysis to identify enriched pathways and to identify the biological significance of the candidate genes, using an FDR cut-off of 0.05.

Biological Process (Fig. 5A) showed that the most enriched term by fold enrichment was regulation of nitrogen utilization (∼780-fold), followed by cGMP-mediated signaling (∼100-fold). Other enriched processes included lymphocyte differentiation, T cell activation, regulation of protein phosphorylation, lymphocyte and leukocyte activation, carbohydrate derivative metabolism, and phosphorylation-related terms. Cell death processes, including regulation of apoptosis, programmed cell death, and negative regulation of signaling, were also enriched, with −log10(FDR) values between 3.5 and 3.7. This points to immune activation, metabolic regulation, and cell death as core shared processes.

**Fig. 5A.**
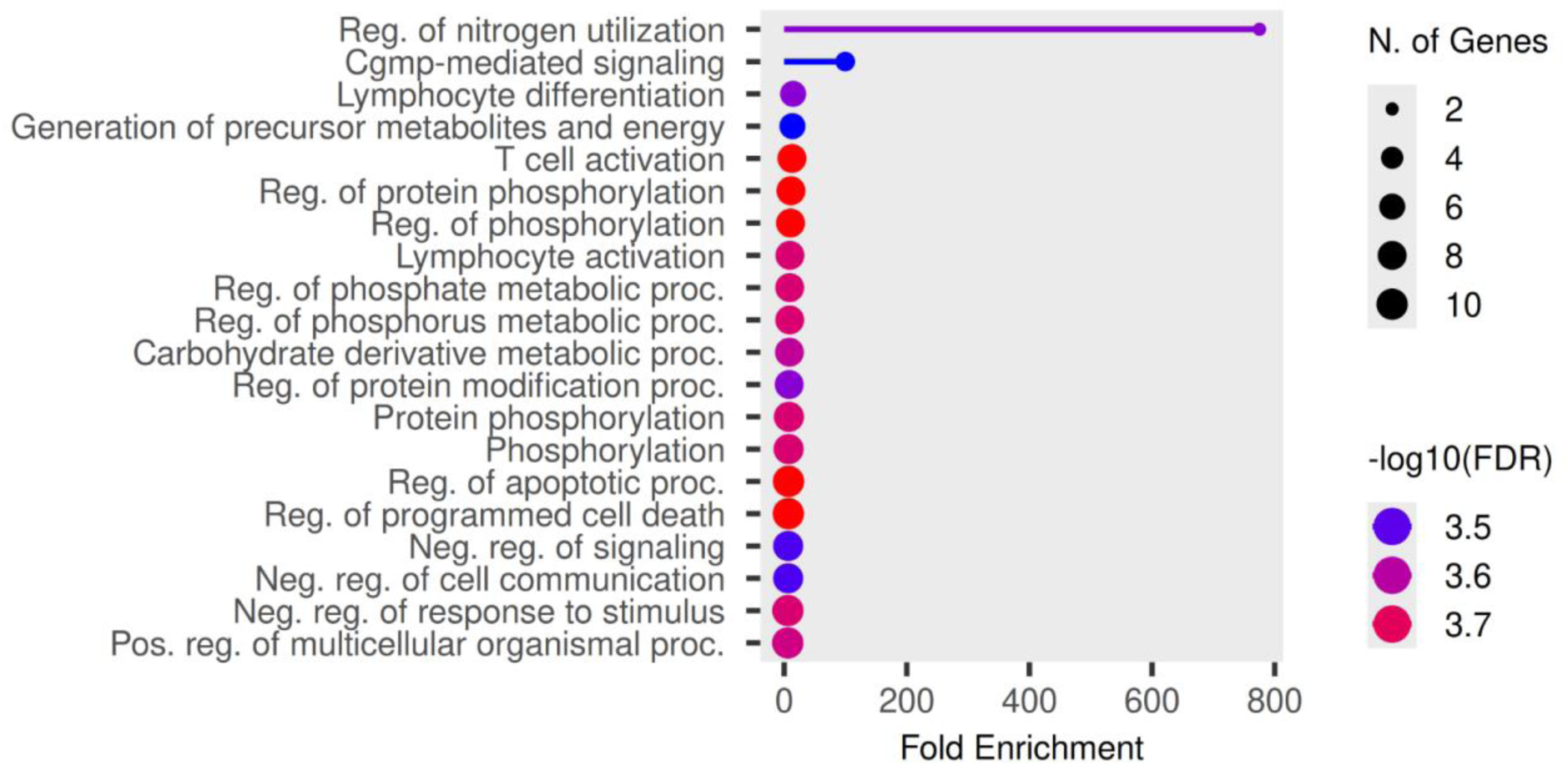
Biological Processes

Cellular Component (Fig. 5B) showed that the Bcl-2 family protein complex was the most enriched term (∼210-fold; −log10(FDR) ≈ 2.6), followed by the pore complex (∼85-fold). Mitochondrial terms, including the mitochondrial outer membrane, organelle outer membrane, outer membrane, mitochondrial membrane, and mitochondrial envelope, were also enriched. This pattern links mitochondria-driven apoptosis to the shared biology of both diseases.

**Fig 5B.**
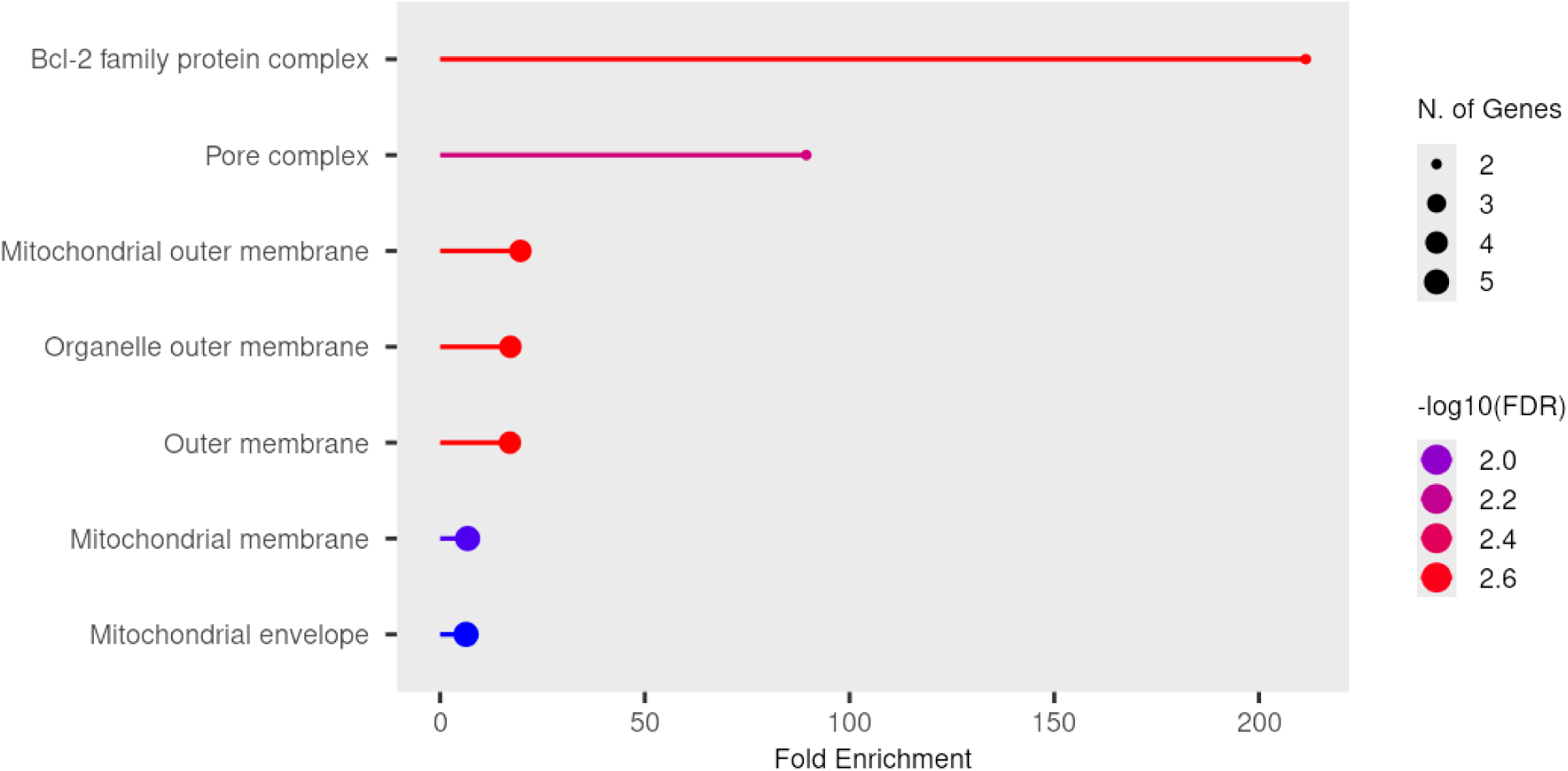
Cellular Processes

Molecular Function (Fig. 5C) showed that the BH3 domain binding was the top term (∼400-fold; −log10(FDR) ≈ 2.4), reflecting the prominence of BCL2-family interactions. RNA polymerase III-related binding terms exceeded 300-fold enrichment, while melatonin receptor activity and leptin receptor binding reached ∼300-fold each. BH domain binding, death domain binding, hormone receptor binding, hormone activity, and ubiquitin-related binding terms were also enriched, highlighting apoptosis, hormonal signaling, and transcriptional regulation as shared molecular functions.

**Fig. 5C.**
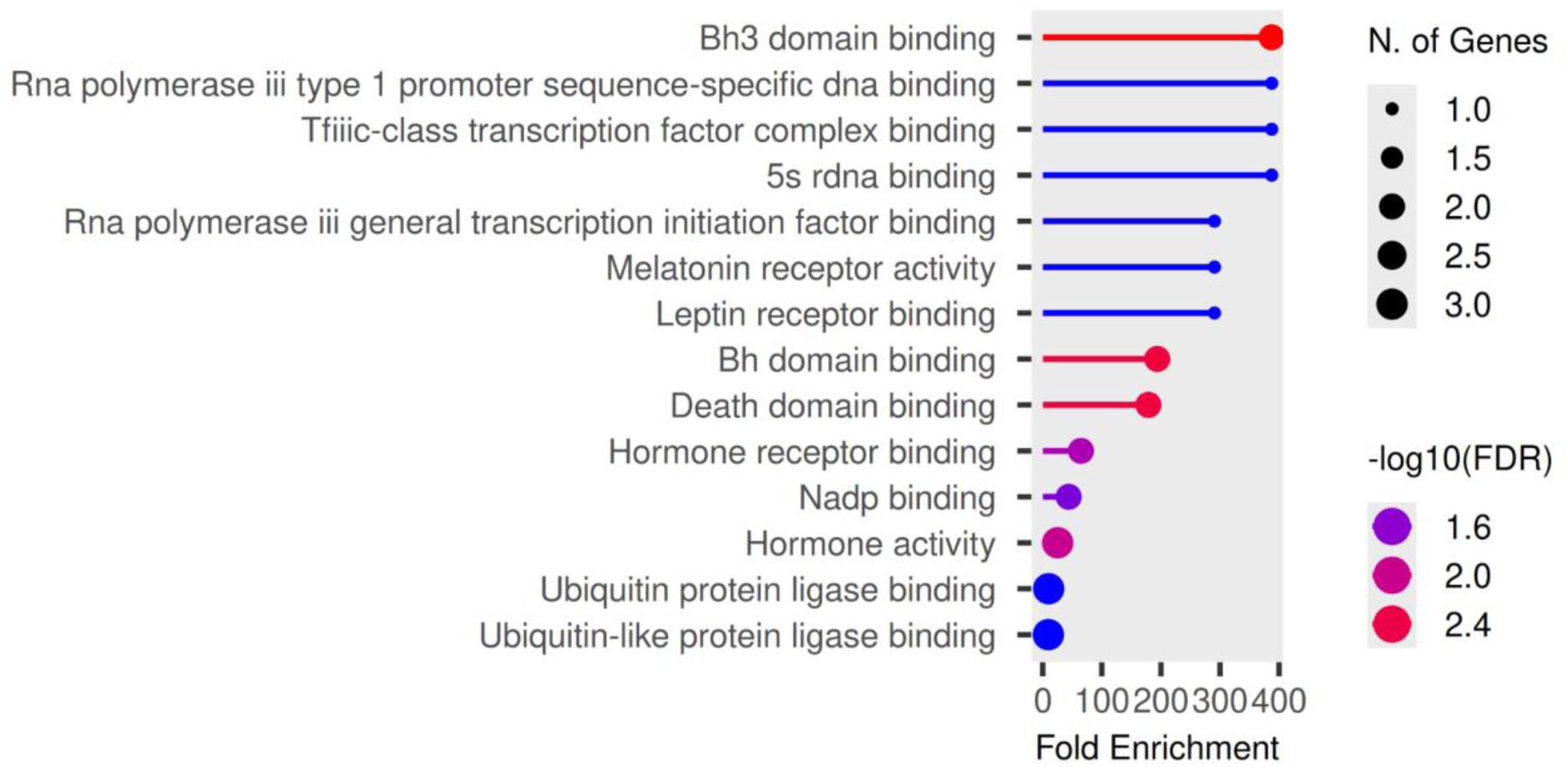
Molecular Function

In the KEGG Pathway (Fig. 5D), three genes were highlighted in red on the pathway map (INS, PIK3, and MTOR). INS appeared at two points in the pathway, reflecting its roles in both insulin receptor activation and pancreatic beta-cell regulation. PIK3 was mapped along the IRS1-PI3K signaling branch, while MTOR appeared within the downstream insulin signaling cascade.

**Fig. 5D.**
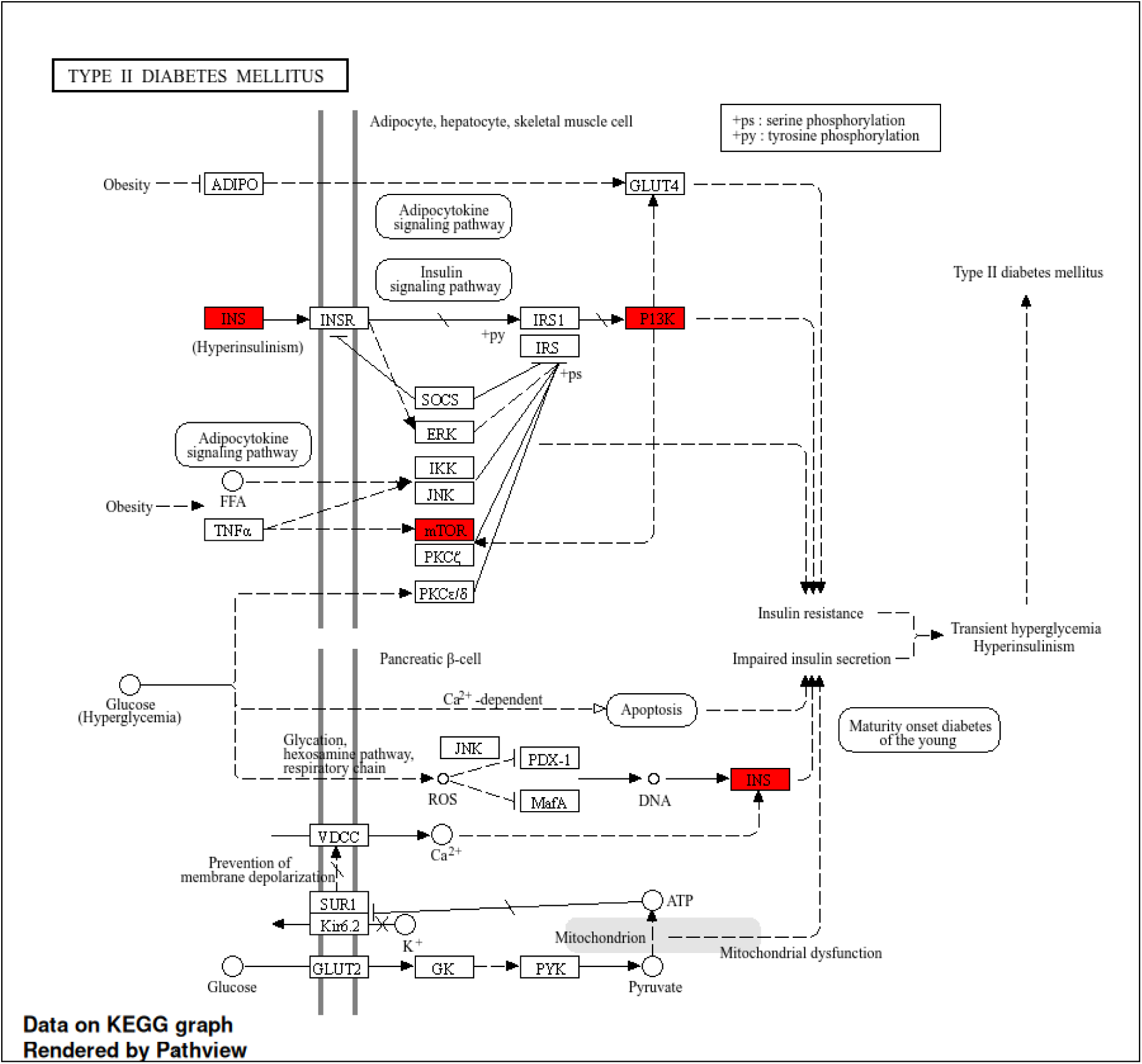
KEGG Pathway

## DISCUSSION

Type 2 diabetes mellitus (T2DM) and polycystic ovary syndrome (PCOS) are among the most prevalent endocrine and metabolic disorders globally, sharing well-established clinical links including insulin resistance, hyperinsulinaemia, obesity, and chronic low-grade inflammation [14, 15]. Despite growing interest in their shared biology, the molecular relationship between them remains incompletely understood. Prior bioinformatics investigations, including the landmark study by Zhang et al., [12] which identified four common genes (BIRC3, DEPTOR, TNNI3, ADRA2A), have focused predominantly on identifying genes shared between both conditions at the transcriptomic level without considering whether those genes are upregulated or downregulated. As Kong et al. [9] demonstrated, confirming whether genes are highly expressed or suppressed is essential. Without this information, shared genes may appear to have similar roles when in fact they act in opposite ways.

This study employed a direction-aware integrative bioinformatics approach to explore the shared molecular mechanisms between T2DM and PCOS. Transcriptomic analysis of GSE25724 (pancreatic islet tissue, T2DM) and GSE138518 (ovarian granulosa tissue, PCOS) identified 1,302 and 225 DEGs, respectively. Despite producing 1,524 DEGs in total, only 3 genes (SLC6A8, RGS4, and SORL1) were shared between the two conditions, and all three were regulated in opposing directions, revealing a pattern of discordance that a simple gene overlap analysis would have entirely missed. To capture the broader genetic connection of both diseases, genes already known to be associated with these diseases were additionally retrieved from CTD, OMIM, and GeneCards, yielding 214 shared disease genes. Combining these with the 3 overlapping DEGs produced a final candidate list of 217 genes, which were taken forward for PPI network construction, hub gene identification, and functional enrichment analysis. PPI network analysis via STRING identified 70 interacting nodes and 64 edges. CytoHubba analysis across six independent algorithms identified 18 high-confidence hub genes, including INS, BCL2, MTOR, LEP, and MFN2, as central molecular players shared between both diseases, with functional enrichment converging on immune activation, mitochondria-driven apoptosis, and insulin signalling as core shared biological themes.

Analysis of the GEO datasets revealed substantially more DEGs in T2DM than in PCOS. In T2DM, this broad transcriptional disruption reflects how pancreatic β-cells are overwhelmed by chronic exposure to high glucose and lipid levels, leading to cellular stress, widespread reprogramming of gene activity, and eventual apoptosis [16, 17]. By contrast, the smaller DEG signature in PCOS indicates that transcriptional changes are more tissue-restricted, occurring primarily within granulosa cells of the ovary rather than across an entire organ such as the pancreas [18]. No genes were concordantly upregulated or downregulated in both diseases simultaneously, a finding that would be entirely missed by a non-directional overlap analysis.

To illustrate how direction-aware analysis reshapes the interpretation, the three overlapping genes, SLC6A8, RGS4, and SORL1, showed opposite regulation in T2DM and PCOS. The creatine transporter SLC6A8 (CT1) was upregulated in T2DM pancreatic islets but downregulated in PCOS granulosa cells. The creatine transport system is central to cellular energy buffering via the creatine–phosphocreatine shuttle, which regenerates ATP under conditions of high metabolic demand [19]. In T2DM, this upregulation likely reflects a compensatory response to the increased energetic demands on chronically stressed beta-cells, given the critical role of the creatine-phosphocreatine shuttle in cellular ATP regeneration [20, 21]. Conversely, its downregulation in PCOS may indicate impaired granulosa cell bioenergetics, reducing the metabolic support required for normal folliculogenesis and oocyte maturation [21, 22].

RGS4 (Regulator of G Protein Signalling 4) showed the opposite pattern, being upregulated in PCOS but downregulated in T2DM. RGS4 functions as a GTPase-activating protein that attenuates G-Protein Coupled Receptor (GPCR) signalling by accelerating GTP hydrolysis on Gα subunits [23]. In PCOS granulosa cells, increased RGS4 expression may represent a compensatory attempt to dampen aberrant GPCR-mediated signalling, particularly through luteinising hormone (LH) and follicle-stimulating hormone (FSH) receptors, which are dysregulated in PCOS [24]. Elevated LH signalling contributes to androgen excess and follicular arrest, and RGS4 upregulation could reflect an attempt to limit this hyperactivation. In T2DM islets, reduced RGS4 may prolong pro-secretory GPCR signalling through incretin receptors such as Glucagon-like peptide-1 receptor (GLP-1R) and Glucose-Dependent Insulinotropic Polypeptide Receptor (GIPR), potentially enhancing insulin secretion [25]. However, sustained activation under chronic stress can exacerbate calcium overload and metabolic exhaustion, accelerating β-cell dysfunction [26].

Finally, SORL1, a multifunctional vesicular sorting receptor [27], was upregulated in PCOS but downregulated in T2DM. In pancreatic β-cells, SORL1 supports insulin granule biogenesis and vesicular trafficking; its downregulation likely contributes to the secretory dysfunction characteristic of T2DM. This role is consistent with evidence from its close structural homologue SORCS1, a VPS10P domain receptor whose loss severely impairs insulin secretory granule biogenesis in beta cells [28, 29]. In PCOS granulosa cells, SORL1 upregulation may reflect enhanced endosomal trafficking in response to hyperandrogenic and hyperinsulinaemic conditions, potentially altering receptor recycling for gonadotrophins or insulin receptors at the cell surface [30]. Together, the discordant regulation of all three genes demonstrates that even shared transcripts are recruited into opposing molecular responses shaped by their distinct tissue environments, a finding that would be overlooked by a gene overlap analysis that ignores directionality.

The striking discrepancy between DEG-level overlap (0.2%) and disease gene-level overlap (21%) reflects a biologically meaningful distinction rather than a methodological inconsistency. DEG analysis captures the immediate transcriptional response of a specific tissue at a particular disease stage, whereas disease gene databases aggregate evidence across GWAS, Mendelian genetics, and biochemical studies, many of which leave no detectable transcriptomic footprint in a single dataset. The 214 shared disease genes therefore represent the deeper genetic infrastructure linking T2DM and PCOS, providing a molecular rationale for the elevated lifetime T2DM risk observed in women with PCOS. Their convergence on the same biological themes identified in hub-gene and enrichment analyses confirms that this overlap is functionally coherent, underscoring shared pathogenic pathways.

Functional enrichment analysis of the 217 candidate genes shared between T2DM and PCOS revealed convergent biological themes spanning immune dysregulation, mitochondrial apoptosis, and dysregulated cell death signaling, revealing the broad multisystem overlap between these conditions. Regulation of nitrogen utilization emerged as the most enriched biological process (∼780-fold), followed by cGMP-mediated signaling (∼100-fold). Although the extreme fold enrichment for nitrogen utilization likely reflects the small denominator effect inherent to ShinyGO’s metric, the biological relevance remains credible, as nitric oxide (NO), a nitrogen-derived signaling molecule is a key vasodilator whose impaired production has been mechanistically linked to insulin resistance and vascular dysfunction in T2DM [31], and growing evidence implicates NO dysregulation in the ovarian microenvironment of PCOS, where insulin resistance drives endothelial dysfunction through the NO pathway [32]. The enrichment of cGMP-mediated signaling is consistent with this, since NO exerts many of its downstream effects via cGMP-dependent pathways [31]. Immune-related processes, including lymphocyte differentiation, T cell activation, and leukocyte activation, were also enriched, reinforcing the well-established role of chronic low-grade inflammation as a shared pathogenic driver in both T2DM and PCOS [33].

At the cellular component level, enrichment of the BCL-2 family protein complex, pore complex, and mitochondrial compartments highlights a mitochondria-centric pattern that is mechanistically coherent: the BCL-2 family governs the intrinsic apoptosis pathway by regulating outer membrane permeabilization [34], a process reported to be dysregulated in pancreatic β-cells in T2DM, where an imbalance between pro-apoptotic proteins (BAD, BAX) and anti-apoptotic members (BCL-2, BCL-xL) drives β-cell loss and chronic hyperglycemia. In PCOS, excessive granulosa cell apoptosis driven by mitochondrial dysfunction, hormonal imbalance, and oxidative stress contributes to follicular atresia and ovarian dysfunction [35], while co-enrichment of the pore complex further implicates mitochondrial permeability transition in the shared pathobiology of both conditions. At the molecular function level, BH3 domain binding was the top-enriched term, consistent with the dominance of BCL-2 family interactions observed in the cellular component analysis, as BH3-only proteins are critical pro-apoptotic mediators that antagonize anti-apoptotic BCL-2 members [34]. These enrichment results showed immune dysregulation, NO/cGMP signaling, and mitochondrial apoptosis as convergent pathways linking T2DM and PCOS, with fold enrichment values interpreted with caution, but biological coherence was strongly supported across multiple levels of analysis.

We identified 18 hub genes in the shared PCOS and T2DM PPI network using CytoHubba across six independent scoring algorithms, suggesting that these genes may be crucial to the overlapping molecular pathobiology of both conditions. Among the most prominent hub genes, INS emerged as the most highly connected node, a finding that is biologically expected given the central role of insulin dysregulation in both diseases [36]. In T2DM, impaired insulin secretion combined with peripheral insulin resistance drives chronic hyperglycaemia, while in PCOS, hyperinsulinaemia directly stimulates ovarian theca cell androgen production by amplifying LH-mediated steroidogenesis and increasing cytochrome P450c17α activity, contributing to the hyperandrogenism that defines the syndrome. The prominence of INS in the shared interaction network positions insulin as the molecular axis around which much of the pathophysiological connection between T2DM and PCOS resolves. Closely linked to INS, PIK3CD encodes the delta catalytic subunit of PI3K, a key mediator of downstream insulin receptor signaling that governs glucose uptake and glycogen synthesis through AKT activation; impairment of this PI3K/AKT pathway is now recognized as central to insulin resistance in both T2DM and PCOS [37]. mTOR further reinforces this cluster; chronic mTORC1 hyperactivation creates a negative feedback loop that suppresses IRS1/2 signalling and worsens insulin resistance in T2DM, while in PCOS, dysregulated mTOR signalling in the ovary has been implicated in aberrant granulosa cell proliferation and follicular development [38, 39]

The co-identification of BCL2 and BAX as hub genes, alongside strong enrichment of BCL2 - family protein complexes and BH3 domain binding as the top cellular component and molecular function terms, respectively, places mitochondria-mediated apoptosis at the centre of the shared pathobiolog [34]. BCL2 and BAX are the archetypal anti- and pro-apoptotic regulators of the intrinsic apoptotic pathway, and their balance governs cellular commitment to apoptosis through mitochondrial outer membrane permeabilisation [40, 41]. In T2DM, increased BAX and decreased BCL2 expression in diabetic islets drives progressive beta-cell loss through glucotoxicity, lipotoxicity, and endoplasmic reticulum stress [42, 43]. In PCOS, an elevated BAX/BCL2 ratio has been directly documented in granulosa cells of polycystic ovaries, where dysregulated apoptosis disrupts normal follicular turnover and contributes to the accumulation of small antral follicles characteristic of the syndrome [44]. The identification of both genes as shared hub genes therefore provides a mechanistic basis for understanding how apoptotic dysregulation, operating through the same mitochondrial pathway, contributes to tissue-specific pathology in both diseases.

LEP (leptin) is a hormone secreted primarily by adipose tissue that regulates energy homeostasis, appetite, and reproductive function through its central and peripheral receptors [45, 46]. Leptin resistance, a state in which the hypothalamus and peripheral tissues fail to respond normally to circulating leptin, contributes to impaired energy balance regulation and exacerbates insulin resistance in T2DM [46]. In PCOS, elevated leptin levels interfere with oocyte maturation and ovarian steroidogenesis, with hyperleptinemia reported independently of body mass index. MFN2 (Mitofusin-2) encodes an outer mitochondrial membrane GTPase essential for mitochondrial fusion, a process that maintains mitochondrial network integrity, bioenergetic efficiency, and quality control [47]. Reduced MFN2 expression has been documented in the skeletal muscle of insulin-resistant individuals and T2DM patients, contributing to mitochondrial fragmentation, impaired oxidative phosphorylation, and worsened insulin signalling. Its identification as a hub gene is further supported by evidence that MFN2 is significantly downregulated in PCOS oocytes and granulosa cells, where disruption of mitochondria-associated ER membrane (MAM) structure impairs calcium transfer and energy metabolism, linking mitochondrial fragmentation to ovarian dysfunction [47].

MTNR1B is one of the most consistently replicated T2DM genetic risk loci identified in genome-wide association studies, with the rs10830963 risk variant associated with impaired early-phase insulin secretion through upregulation of MT2 receptors in pancreatic beta cells and enhanced melatonin-mediated inhibition of adenylate cyclase. Its identification as a shared hub gene in the PCOS-T2DM network is particularly notable, as MTNR1B variants have also been directly linked to PCOS risk, and melatonin receptors expressed on ovarian granulosa cells have been implicated in oocyte maturation and follicular function, providing a direct mechanistic connection between circadian signalling dysregulation and the reproductive-metabolic dysfunction shared by both conditions [47] NPPA (natriuretic peptide A), encoding atrial natriuretic peptide (ANP), has emerging roles in metabolic regulation beyond its classical functions in blood pressure and fluid homeostasis [48, 49]. ANP has been shown to stimulate lipolysis in adipocytes, influence insulin sensitivity [49], and regulate mitochondrial biogenesis through cGMP signalling [50], consistent with the enrichment of cGMP-mediated signalling as the second most enriched biological process term in this study. The identification of NPPA as a hub gene and the enrichment of cGMP signalling suggest that natriuretic peptide-cGMP pathways may represent an underappreciated shared mechanism in T2DM and PCOS, particularly in the context of cardiovascular and metabolic comorbidity that both diseases share [51].

Several additional hub genes show further mechanistic dimensions. MTHFR dysfunction leads to hyperhomocysteinaemia, which has been associated with insulin resistance and endothelial dysfunction in T2DM [51], and elevated homocysteine levels have been documented in PCOS patients, where they may contribute to the cardiovascular risk accompanying the syndrome. H6PD regulates intracellular glucocorticoid activation through its control of 11β-HSD1 activity in the endoplasmic reticulum; dysregulation of this axis contributes to visceral obesity and insulin resistance common to both diseases, with altered glucocorticoid metabolism also proposed as a contributor to adrenal androgen excess in PCOS. TARDBP and PRDM16 represent emerging areas of mechanistic interest; TARDBP has reported roles in RNA metabolism under metabolic stress conditions, while PRDM16, a master regulator of brown adipocyte identity, raises the possibility that thermogenic adipose biology intersects with the shared pathophysiology of both diseases, an area warranting dedicated experimental investigation [52]. The remaining hub genes GNB1, TNFRSF9, TNFRSF14, EXOSC10, and ATAD3A were consistently identified across all six CytoHubba algorithms and represent candidates for prioritisation in future experimental validation studies.

KEGG pathway enrichment analysis identified the Type II Diabetes Mellitus pathway as a central signaling network linking T2DM and PCOS, integrating multiple biological processes including insulin resistance, inflammatory signaling, mitochondrial dysfunction, and β-cell apoptosis [53]. The pathway illustrates how metabolic stress disrupts insulin receptor signaling through IRS– PI3K-mediated cascades, ultimately impairing glucose uptake and promoting chronic hyperglycemia. Under physiological conditions, insulin binding to the insulin receptor activates IRS proteins and PI3K signaling, which promotes GLUT4 translocation and cellular glucose uptake [53]. However, chronic metabolic overload associated with obesity and hyperinsulinemia disrupts this pathway through inhibitory signaling mechanisms. A major feature of the KEGG pathway is the involvement of inflammatory mediators and stress-responsive kinases, including TNF-α, JNK, IKK, and ERK. These signaling molecules impair insulin receptor substrate (IRS) activity through inhibitory serine phosphorylation, thereby reducing downstream insulin signaling efficiency and promoting insulin resistance [54]. Chronic low-grade inflammation has been widely implicated in both T2DM and PCOS, where elevated inflammatory cytokines contribute to metabolic dysfunction, endocrine imbalance, and impaired insulin sensitivity [53]. The enrichment of inflammatory signaling within the KEGG pathway therefore reinforces the immune-related biological processes identified in the present study.

The pathway also highlights PI3K signaling, which emerged as a key node within the insulin signaling cascade. PI3K regulates glucose metabolism through downstream activation of pathways controlling glucose transporter translocation and energy utilization [55]. Dysregulation of PI3K signaling is strongly associated with impaired insulin action and metabolic dysfunction [55]. This observation is particularly relevant given the identification of PIK3CD as a hub gene in the present analysis, supporting its role as a shared mediator of insulin resistance between T2DM and PCOS. Another important component of the pathway is mTOR, which functions as a nutrient-sensing regulator of growth and metabolism. Persistent activation of mTOR signaling contributes to negative feedback inhibition of IRS proteins, worsening insulin resistance. This mechanistic relationship supports the identification of MTOR as a hub gene and suggests that excessive nutrient signaling may represent a common pathogenic mechanism shared across metabolic and reproductive dysfunction [56].

At the pancreatic β-cell level, the KEGG pathway demonstrates how chronic hyperglycemia and oxidative stress promote mitochondrial dysfunction, calcium imbalance, and apoptosis. β-cell apoptosis is recognized as a major contributor to T2DM progression, as loss of insulin-producing cells limits the ability to compensate for peripheral insulin resistance [56]. Similar apoptotic mechanisms have also been implicated in PCOS granulosa cells, where oxidative stress and mitochondrial dysregulation contribute to follicular dysfunction [57]. These findings align closely with the enrichment of mitochondrial apoptosis pathways observed in the current study. The KEGG pathway provides mechanistic support for the identified hub genes and enrichment results, illustrating how insulin resistance, inflammation, mitochondrial dysfunction, and apoptotic signaling converge to drive disease progression [58].

## CONCLUSION

This study used an integrative bioinformatics approach to evaluate the molecular relationship between PCOS and T2DM by combining differential gene expression, curated disease-gene databases, protein–protein interaction network analysis, hub gene identification, and functional enrichment analysis. Although only three DEGs were shared between the two conditions, all showed discordant expression patterns, indicating distinct transcriptional responses. In contrast, the substantial overlap of disease-associated genes and the identification of 18 high-confidence hub genes revealed common molecular networks enriched in insulin signaling, apoptosis, mitochondrial function, immune regulation, and metabolic processes. These findings provide a more comprehensive understanding of the molecular links between PCOS and T2DM and identify potential candidate genes and pathways for future mechanistic and therapeutic studies.

## Supporting information

references

## LIST OF ABBREVIATIONS

PCOS: Polycystic ovarian syndrome
T2DM: Type 2 Diabetes Mellitus
PPI: Protein-protein Interaction
DEGs: differentially expressed genes
NCBI: National Center for Biotechnology Information
GEO: Gene Expression Omnibus
GSE: Gene series accessions
GEO2R: Gene Expression Omnibus 2 R
CTD: Comparative Toxicogenomics Database
OMIM: Online Mendelian Inheritance in Man
STRING: Search Tool for the Retrieval of Interacting Genes
DAGs: Directed acyclic graphs
MCC: Maximal Clique Centrality
MNC: Maximum Neighborhood Component
KEGG: Kyoto Encyclopedia of Genes and Genomes
GO: Gene Ontology
FDR: Functional Enrichment Analysis
RGS4: Regulator of G Protein Signalling 4
GPCR-G: Protein Coupled Receptor
LH: Luteinising Hormone
FSH: Follicle-Stimulating Hormone
GLP-1R: Glucagon-like peptide-1 receptor
GIPR: Glucose-Dependent Insulinotropic Polypeptide Receptor
NO: Nitric Oxide

## DECLARATIONS

### Consent for publication

Not applicable

### Competing interest policy

The authors declare that they have no competing interests.

### Authors’ Contribution

**Abdulsalam Sodiq Okikiola**: Conceptualization, study design, experimental procedures, manuscript drafting, manuscript writing (introduction), and manuscript revision.

**Merylin Wuraola Ogunlola**: Experimental procedures, data collection and analysis, manuscript writing (methodology), and manuscript revision.

**Damilare Alabi Akanbi**: Data validation, manuscript writing (background), and manuscript revision.

**Temiloluwa Esther Aloba**: Data collection, data interpretation, manuscript writing (methodology), and manuscript revision.

**David Ifeoluwa Owopetu**: Study supervision, project administration, manuscript writing (discussion), and critical review of the manuscript.

All authors contributed to the conduct of the experiments, participated in the preparation and revision of the manuscript, and approved the final version of the manuscript.

## Acknowledgements

Not applicable

